# Secondary osteons scale allometrically in mammalian humerus and femur

**DOI:** 10.1101/131300

**Authors:** A. A. Felder, C. Phillips, H. Cornish, M. Cooke, J. R. Hutchinson, M. Doube

## Abstract

Intra-cortical bone remodelling is a cell-driven process that replaces existing bone tissue with new bone tissue in the bone cortex, leaving behind histological features called secondary osteons. While the scaling of bone dimensions on a macroscopic scale is well known, less is known about how the spatial dimensions of secondary osteons vary in relation to the adult body size of the species. We measured the cross-sectional area of individual intact secondary osteons and their central Haversian canals in transverse sections from 40 stylopodal bones of 39 mammalian species. Scaling analysis of our data shows that mean osteonal resorption area (negative allometry, exponent 0.23, *R*^2^ 0.54, *p* < 0.005) and Haversian canal area (negative allometry, exponent 0.34, *R*^2^ 0.45, *p* < 0.005) are significantly related to body mass, independent of phylogeny. This study is the most comprehensive of its kind to date, and allows us to describe overall trends in the scaling behaviour of secondary osteon dimensions, supporting the inference that osteonal resorption area may be limited by the need to avoid fracture in smaller mammalian species, but the need to maintain osteocyte viability in larger mammalian species.

## 1. Introduction

The cell-driven process of replacing existing cortical bone tissue with new packets of osteoid (a collagenous matrix which later mineralisses, becoming bone) is termed intracortical bone remodelling and involves the coordinated action of osteoclasts (bone resorbing cells) and osteoblasts (osteoid forming cells) around a central blood vessel, contained within the Haversian canal. Some osteoblasts are buried in the osteoid, becoming osteocytes, but remain connected to each other and to the Haversian canal via the lacunocanalicular network. Secondary osteons, the remnants of such remodelling events, are approximately cylindrical features at a sub-millimetre scale. In the diaphyses of long bones, osteons are typically oriented in an approximately longitudinal direction [1,2]. They enclose the Haversian canal and are delineated by a collagen-deficient border [3], the cement sheath. Haversian canals form a complex vascular network with branching of different types [4].

The function of intra-cortical bone remodelling is the subject of much discussion in the literature. Several mechanical and metabolic functions have been put forward, which are not mutually exclusive. The most widespread hypothesis of bone remodelling function is that bone tissue accumulates micro-damage through everyday loading, and therefore needs to be continuously replaced throughout the lifetime of an organism [5] to avoid fatigue fracture. Although this idea is coherent with micro-damage accumulation in cyclically-loaded, synthetic materials, bone remodelling does not seem always to occur in regions of high stress, where the most micro-damage would be expected [6]. This may be because the infilling and mineralisation process takes several months to complete (in humans) [7], and thus the presence of large resorption cavities may temporarily weaken highly stressed regions, as well as incurring considerable metabolic costs [6]. Other mechanical functions may be that the cement sheath deflects cracks into less critical directions, thus enhancing toughness by dissipating energy [8] and protecting the nerves, blood vessels and osteocytes within the secondary osteons [9]. The collagen fibre orientation can vary between adjacent lamellae deposited within the resorbed area when the osteon is forming, giving rise to typical osteon “morphotypes” [10,11], which may be adapted to the loading mode [12]. Apart from its roles in maintaining structural integrity, intra-cortical bone remodelling may also regulate calcium homeostasis [13] and mineralisation levels [14], maintain the viability of osteocytes [15], or process the metabolite flow to ensure “intended” limb proportions during growth [16].

In contrast to the generous attention that scaling relationships of whole bones (e.g. [17–19]) and, more recently, of trabeculae [20–22] have received, quantitative measurements of the spatial dimensions of secondary osteons across a large sample of species are yet to be reported in a single study. Furthermore, the presence or absence of a scaling relationship between body mass and secondary osteon dimensions has yet to be fully established. In mammals, secondary osteons are typically found only in species of adult mass > 2 kg, although they can be induced by supra-physiological levels of exercise in rats [23], and tend to occur more extensively in larger species [24]. Secondary osteons are also found within the trabeculae of larger species, including some primates [21]. This study, however, focuses on secondary osteons in cortical bone only. A comparative study of secondary osteon dimensions was performed by Jowsey [25]. The data from this study were later re-used, in combination with data from a study by Tarach and Czaja [26], to investigate scaling relationships between osteonal dimensions and animal size [27]. These studies concluded that osteon size increases with body size for species < 10 kg, and then plateaus. This remains the only such study we are aware of, is based on a small sample of 9 species and does not take into account phylogenetic effects [28]. The spatial dimensions of secondary osteons and Haversian canals have previously been measured for humans [29] and various other mammalian species (e.g. [30–36]) without considering possible inter-specific scaling effects. Histomorphometrical measurements of secondary osteons remain unreported for most mammalian species. Consequently, a broad overview of scaling of secondary osteon dimensions, able to elucidate overall trends that may be representative of the bio-physical constraints within which the intra-cortical bone remodelling process operates, is still missing.

In this study, we address this shortcoming. We seek to establish the parameters of any scaling relationships between the dimensions of the intra-cortical bone remodelling process and animal size. In order to achieve this, we first generated composite images of transverse histological sections of mammalian femora and humeri from a museum collection. We then hand-traced the borders of all intact secondary osteons and their Haversian canals in these images. Finally, we measured their areas and analysed these measurements using the standardized reduced major axis method [37] and phylogenetically independent contrasts [28].

## 2. Materials and Methods

### (a) Specimens

We obtained access to the historical Bd series of John Thomas Quekett’s collection of microscope slides, which today is part of museum collections at the Royal College of Surgeons (RCS) [38]. Quekett’s collection is one of the earliest museum collections for comparative histology of bone. Sections of larger species covered only part of the whole bone cross-section (Figure 1). We selected transverse sections from femur or humerus of mammalian species (total 61 specimens, 56 species, see Tables 1 (humerus) and 2 (femur) for a list of specimens with searchable identifiers). The rightmost columns specify whether any secondary osteons were found. Our further analyses dealt solely with specimens that had secondary osteons. For each species, we also list an estimate for adult body mass, obtained from the PanTHERIA database [39] for extant species and from the study by Larramendi [40] for fossil proboscideans, which we used as an input for our scaling analyses. We chose stylopodal bones, as we expected the least variation in loading patterns among species (although this variation may still be considerable) and the greatest number of secondary osteons in these bones [24].

**Figure 1.**
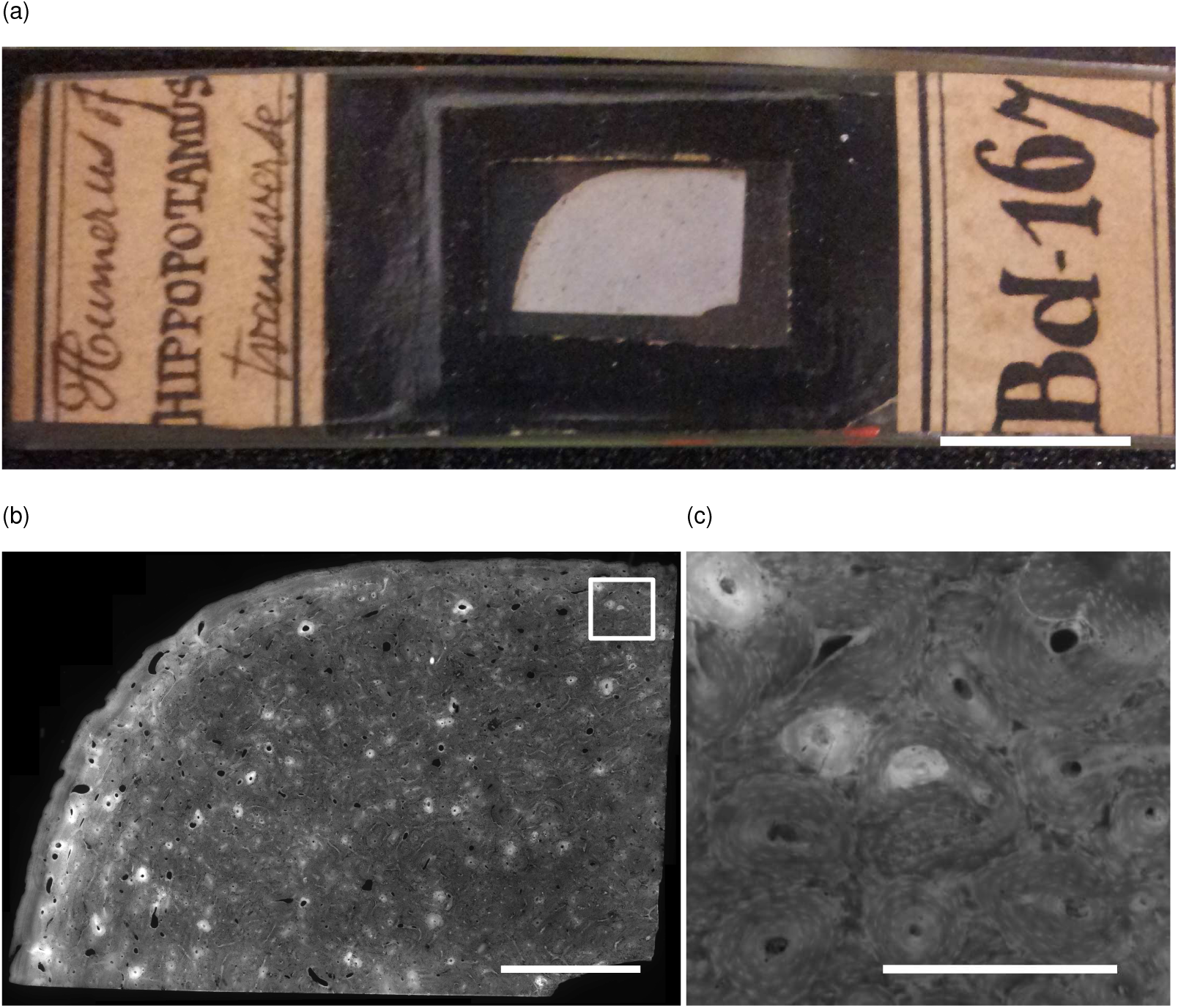
(a) Digital photograph of slide Bd 167 from the Quekett collection (scalebar 10 mm). (b) Fluorescence microscopy image of the same specimen, automatically stitched from 76 manually captured tile images (scalebar 2.5 mm). (c) A close-up of the aforementioned image showing multiple secondary osteons (scalebar 0.5 mm). Images ©The Royal College of Surgeons of England, reproduced here with their kind permission.

**Table 1.**
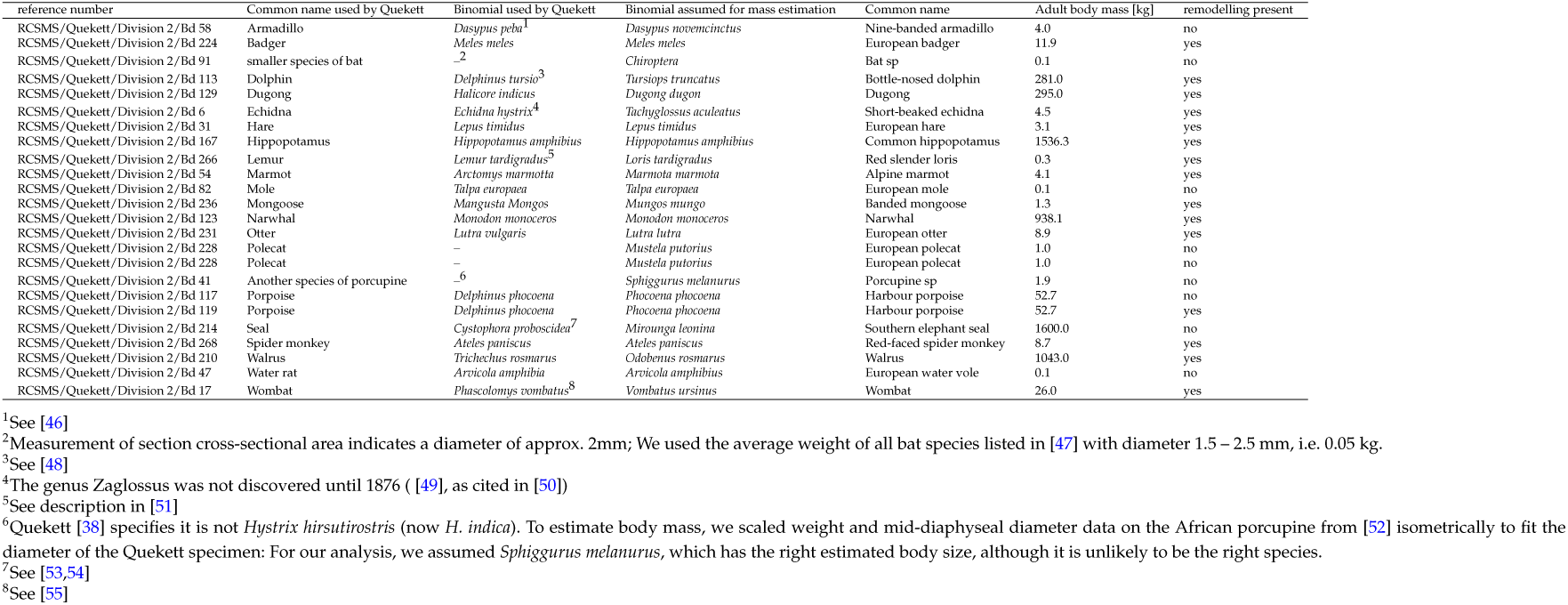
For each humeral specimen: RCS reference number (searchable on the RCS online catalogue SurgiCat), Quekett’s [38] and current terminology, adult body mass estimate [39,40] and whether there were any secondary osteons in it. Secondary literature supporting our assumptions for the terminology can be found in the footnotes.

In some cases, the species was unclear, due to missing or antiquated binomial or common names in the historical documentation [38] of the slides. When this occurred, we usually deduced the taxonomy from the description in the catalogue entry. For some species, we additionally resorted to secondary literature (See footnotes of Tables 1 and 2) to establish a species with more confidence. Out of the six more markedly uncertain cases, two (Bd 137, “fossil elephant” and Bd 139, *Mastodon sp*.) had secondary osteons and were therefore relevant to our main analysis. Analysis of these specimens, as they were fossils, was additionally complicated by possible diagenetic effects. However, scaling parameter estimates calculated excluding these specimens remained comfortably within the overall confidence intervals throughout. Thus, excluding these specimens would not have altered our conclusions.

**Table 2.**
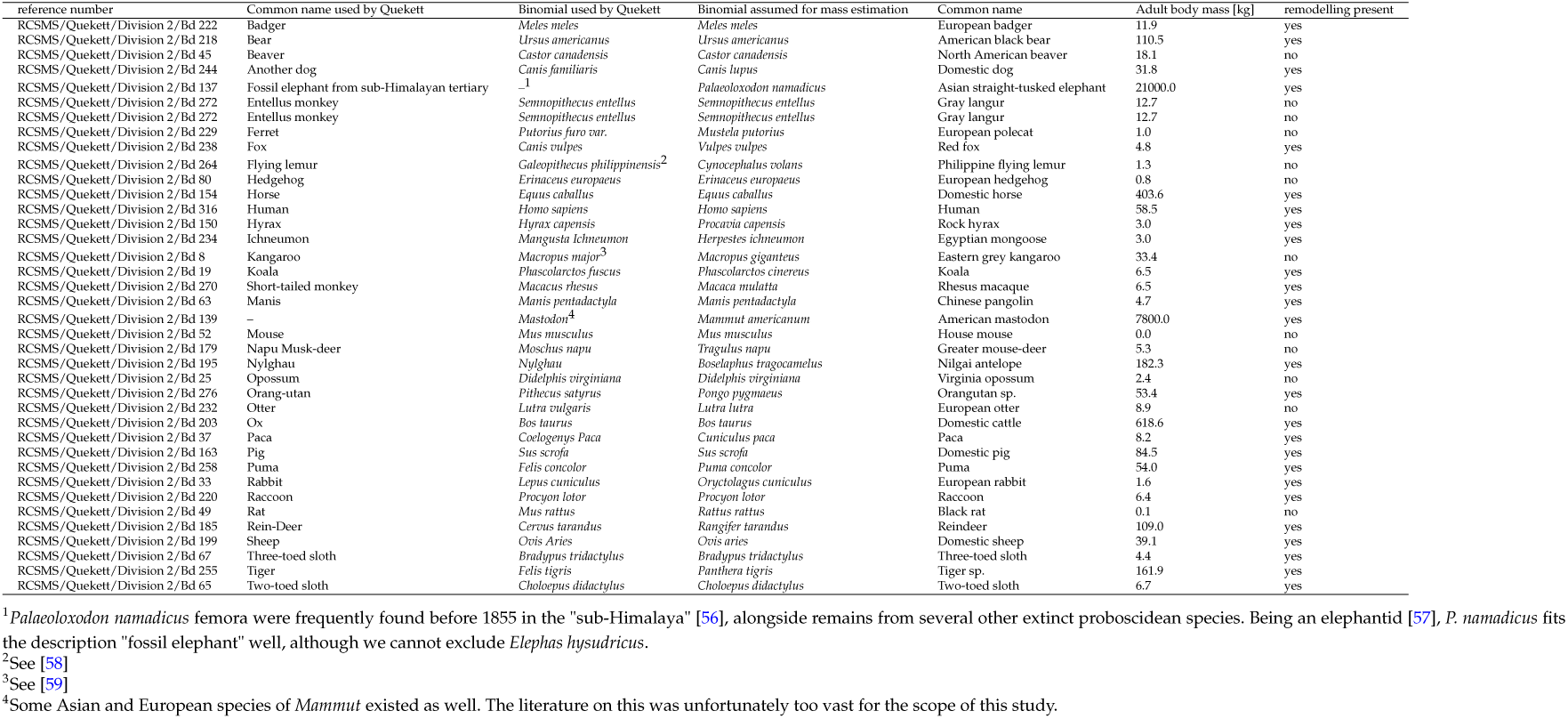
For each femoral specimen: RCS reference number (searchable on the RCS online catalogue SurgiCat), Quekett’s [38] and current terminology, adult body mass estimate [39,40] and whether there were any secondary osteons in it. Secondary literature supporting our assumptions for the terminology can be found in the footnotes.

### (b) Historical details on specimen origin and preparation

The specimens we imaged are likely to have been prepared in different ways by several people. Some display commercial cover slip paper of the time, and may have been purchased by Quekett, while others are likely to have been produced in-house by Quekett or his collaborators. In his treatise on the use of the microscope, Quekett describes dry and wet preparation methods for thin sections of bone, and examples of both are present in the collection today [41]. Exact anatomical location of our samples is unknown, although most seem to be taken along the diaphysis, based on seeing smooth endosteal surfaces with no trabeculae, except, as expected, in the two sloth species [42] and some of the marine mammals [43].

The origin of the samples used in this study is unclear. Some are likely to have originated from the Zoological Society of London (ZSL), London Zoo, as Richard Owen, for whom Quekett was initially an assistant, before becoming his successor, had the right to claim any freshly deceased animal there. Others may have come from skeletons donated to the RCS by John Gould and other 19^th^ century travelling naturalists. Such donations are described by Quekett in his diaries (e.g. entries of 1. October 1840 and 1. December 1842, accessed at RCS library (MS0027/1)), and sections may have been prepared for the microscope later. Another possible source of specimens is the ZSL Osteological Museum, suggested by a letter from 1840 by ZSL honorary secretary William Ogilby offering this collection to the Hunterian museum (RCS-MUS/11/1/14, item 52).

### (c) Imaging

We imaged the specimens in reflected light fluorescence microscopy using a Leitz Laborlux 12 microscope with a built-in LED light (Excitation 450 −490 nm). We manually took overlapping 16-bit gray-scale images, each 1024 by 1024 pixels, covering the entire specimen (4x magnification, 0.12 numerical aperture, pixel size 1.64 μm) with an Orca FLASH4.0LT C11440 digital camera (Hamamatsu Photonics, Hamamatsu City, Japan) and F56-L019 filterset (AHF Analysentechnik, Tübingen, Germany). Consequently, the contrast in the images was generated by green autofluorescence induced by a blue light. In total, we took 3416 images, resulting in an average of 58 image tiles/specimen (minimum 6 (specimen number Bd 91), maximum 180 (Bd 123)). We composed images of entire specimens by applying the Grid/Collections stitching plug-in ([44], version 1.2) of the open-source image processing software Fiji ([45], version 1.51) to the image tiles.

We found intact secondary osteons in 40 specimens (14 humeri, 26 femora) from 39 species. Using a Wacom digitizer (Wacom, Saitama, Japan), we manually traced the boundary of intact secondary osteons and their Haversian canals in the images. Following previously established methods [14], a feature was classified as an intact secondary osteon if >90% of the cement line and the entire canal boundary were visible, and the aspect ratio (maximum diameter/minimum diameter) was < 2, to prevent atypical osteon variants confounding our area measurements. Drifting osteons [60] were included if we could draw a trace around the osteon from the non-drifting side of the cement line along a clear lamellar boundary, and excluded otherwise, as previously done in [14]. We found a fracture callus in specimen Bd 270, which we excluded from analysis. We stored the osteon area (the area within the cement line, which is the cross-section of the cement sheath), the canal area (the area enclosing the Haversian canal) and the infill area (the difference between osteon area and canal area) for each osteon using a custom ImageJ macro (Figure 2). From these directly measured quantities, we also computed two derived quantities for each intact secondary osteon: the infill distance (the distance between the canal border and the cement line, assuming perfectly circular osteons of equivalent area with concentric Haversian canals of equivalent area), and the infill ratio (infill area divided by its corresponding osteon area) using custom ImageJ macros.

**Figure 2.**
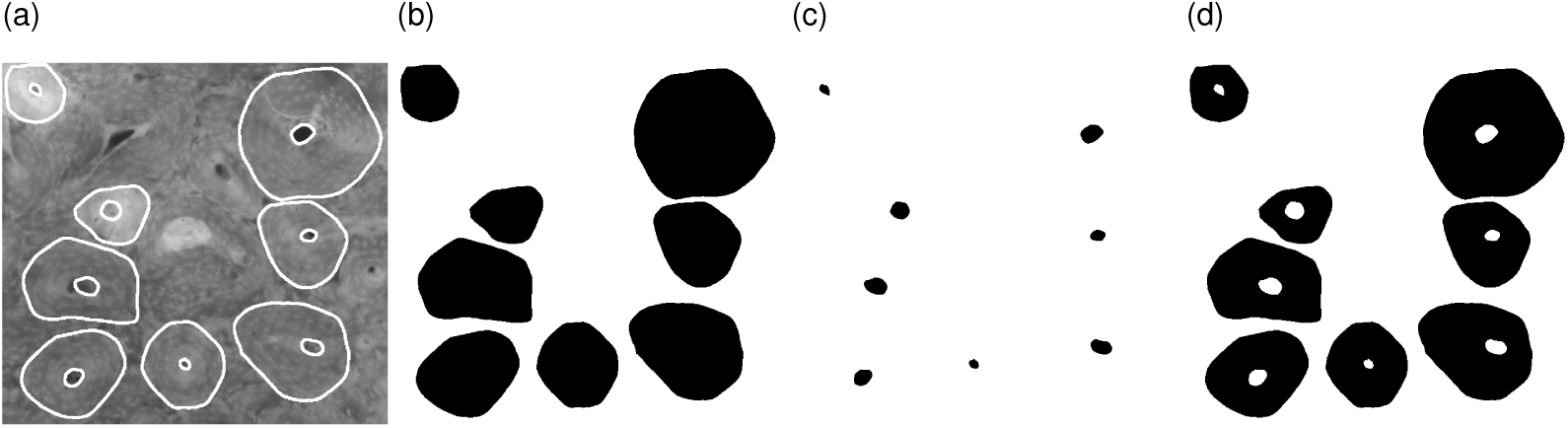
Visualisation of measured data. (a) Close-up of some of the intact secondary osteons and their Haversian canals from a hippopotamus humerus (Bd 167), traced in white (image ©The Royal College of Surgeons of England, reproduced here by their kind permission) (scale: see Figure 1. (c)). (b) Binary file created to measure the osteon areas. (c) Binary file created to measure the canal areas. (d) Binary file created by subtracting the previous binary files from each other to measure the infill areas.

**Figure 3.**
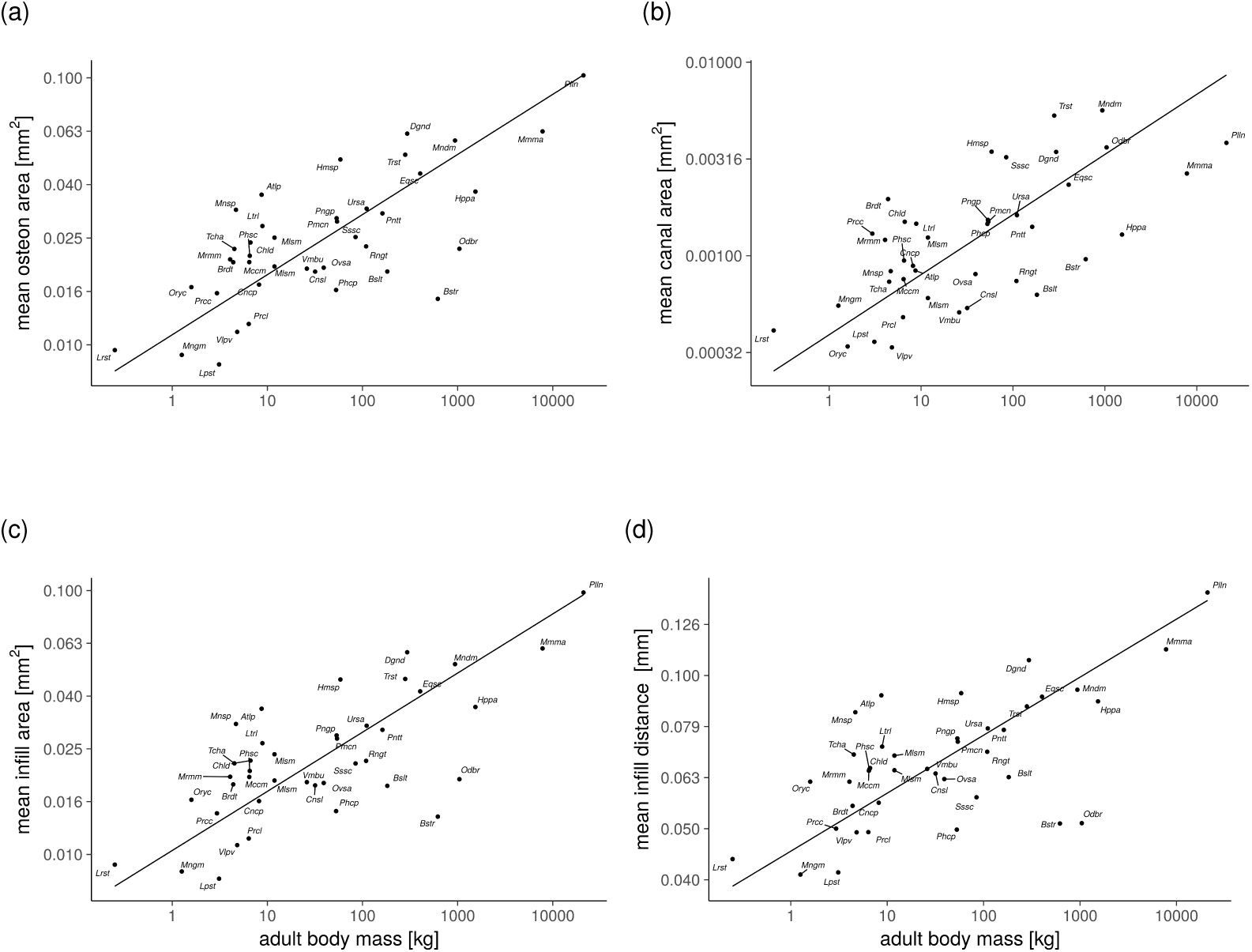
Log-log plots of per species means of the secondary osteon cross-sectional parameters and estimated body mass, where we found a significant relationship (*R*^2^ > 0.3, *p* < 0.005). Estimates, sample square correlation coefficients and *p*-values for all parameters we measured can be found in Tables 3 and 4.

The number of secondary osteons present in each specimen varied from 1 to > 200. To prevent our results from being biased by specimens with low numbers of osteons (often small species, some artiodactyls or marine mammals), we repeated our analysis excluding specimens with fewer than 10 measured osteons. This offered some additional robustness to our analyses, since any false positives (features wrongly classified as secondary) would have a particularly high weight in specimens with few osteons. Again, the results for scaling exponents and elevations were within the bounds obtained including all specimens.

### (d) Statistical analyses

We assumed our quantities of interest *Y* obeyed a power law *Y* = *aM*^*b*^, where *M* denotes body mass. We obtained estimates, lower and upper bounds for both the scaling exponent *b* and the elevation *a* using standardised (reduced) major axis from the R ([61], version 3.3.2) package SMATR 3 [62] on the base-10 logarithms of our variables of interest. We used the same software to test for the null hypotheses of isometry (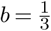 for infill distance, b = 0 for the infill ratio, 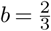 otherwise) and no correlation. We calculated phylogenetically independent contrasts [28] using the `pic` function of the R package `ape` [63] and a species-level phylogenetic tree, including branch lengths [64]. Entries for the extinct proboscideans in our sample were added with a custom R script, using estimated branch lengths from a previous study [65]. SMATR was used to estimate the scaling exponent of independent contrasts, while enforcing zero elevation [37]. In all our standardised major axis analyses, we used *p* < 0.005 and *p* < 0.05 as cut-off values for high and mild statistical significance, respectively. We checked whether the data was normally distributed by incorporating Shapiro-Wilk tests into our code. If normality could not be assumed, we report Spearman’s *ρ*^2^ as a measure of the monotonic relationship between the variables. In the other cases, we report Pearson’s *R*^2^. We interpreted significant relationships with squared (Pearson’s *R* or Spearman’s *ρ*, depending on normality assumptions) correlation coefficients > 0.3 as strong, and otherwise as weak. Our R code [66] and ImageJ macros [67] are available online.

## 3. Results

All logarithmic transformations of our data could be assumed to be normally distributed (Shapiro-Wilk tests, p>0.05), unless we explicitly state this. Also, if scaling exponent and elevation estimates calculated for femora and humeri individually were not comfortably within 95% confidence intervals of the estimates of combined stylopod data, we report the data separately. Otherwise, we pooled the data for analysis.

### (a) Direct measurements

Mean osteon area, canal area and infill area showed negative allometric scaling (*b* = 0.23, *b* = 0.31 and b = 0.22, respectively). These relationships were different to both null hypotheses (scaling-invariance and isometry) with high significance (*p* < 0.005). Sample squared Pearson’s correlation coefficients (*R*^2^) showed that our fits for the means of directly measured variables, where highly significant, explained between 45 and 52% of the variation in our data. The maxima for these quantities showed the same gross pattern as the means, with similar values for *p* and *R*^2^.

Maximum canal area scaled close to isometry, but was still negatively allometric with mild significance (*p* < 0.05). The variation in our data for minimum areas was considerably less well explained by our fits. The correlation of minimum osteon area and minimum infill area with body mass was weak (*R*^2^ = 0.14 and *R*^2^ = 0.14, respectively, *p* < 0.05), and was lost completely when excluding samples that had fewer than 10 osteons (*p* = 0.37). Minimum canal area was not normally distributed (*p* < 0.005, Shapiro-Wilk), and a Spearman’s rank correlation test with body mass yielded a similarly weak relationship (*ρ*^2^ = 0.21, *p* < 0.005, see Figure 5). The elevation for mean canal area was 390 μm^2^, two orders of magnitude less than its analogues for osteon area and infill area, which corresponds to a mean canal diameter of 11.1 μm (assuming circular canals). Estimates and 95% confidence intervals for a and b of directly measured quantities are summarized in Table 3.

**Figure 4.**
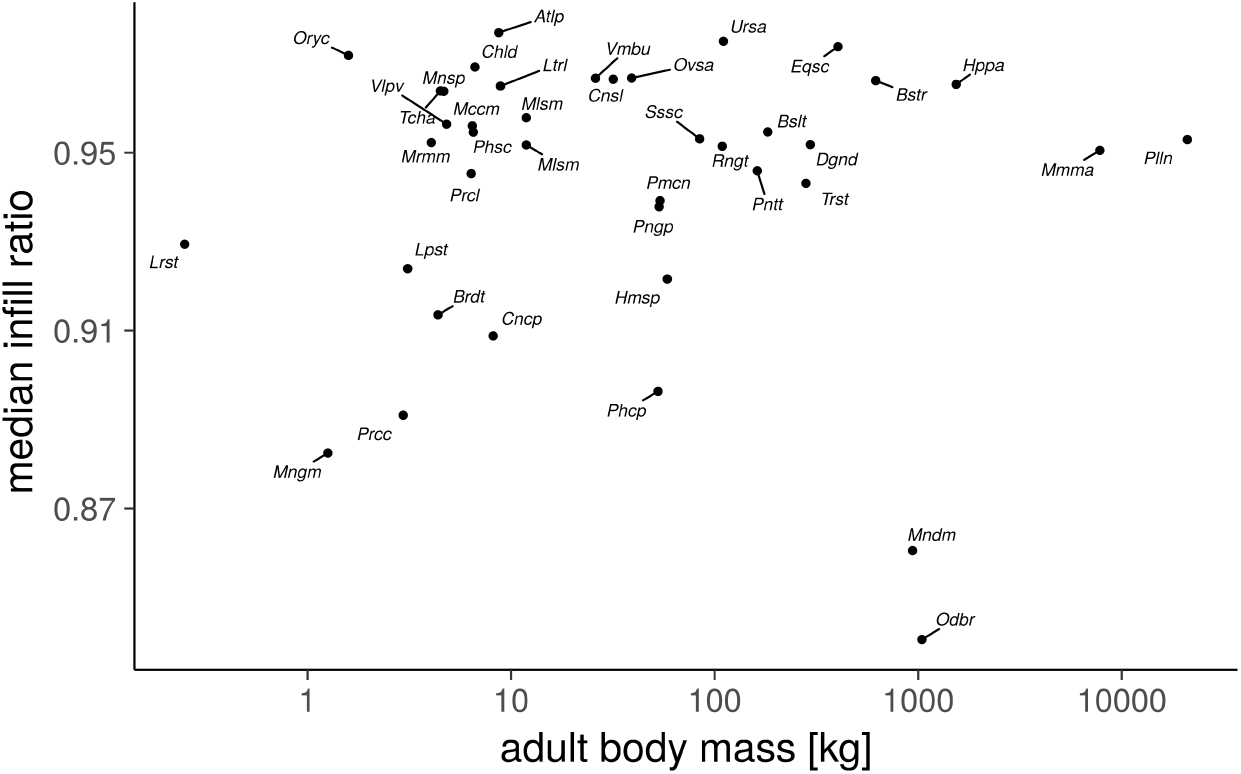
Log-log plot of per species median infill ratio and estimated body mass. Infill ratio does not scale with body mass, and is kept > 85% throughout mammalian species.

**Figure 5.**
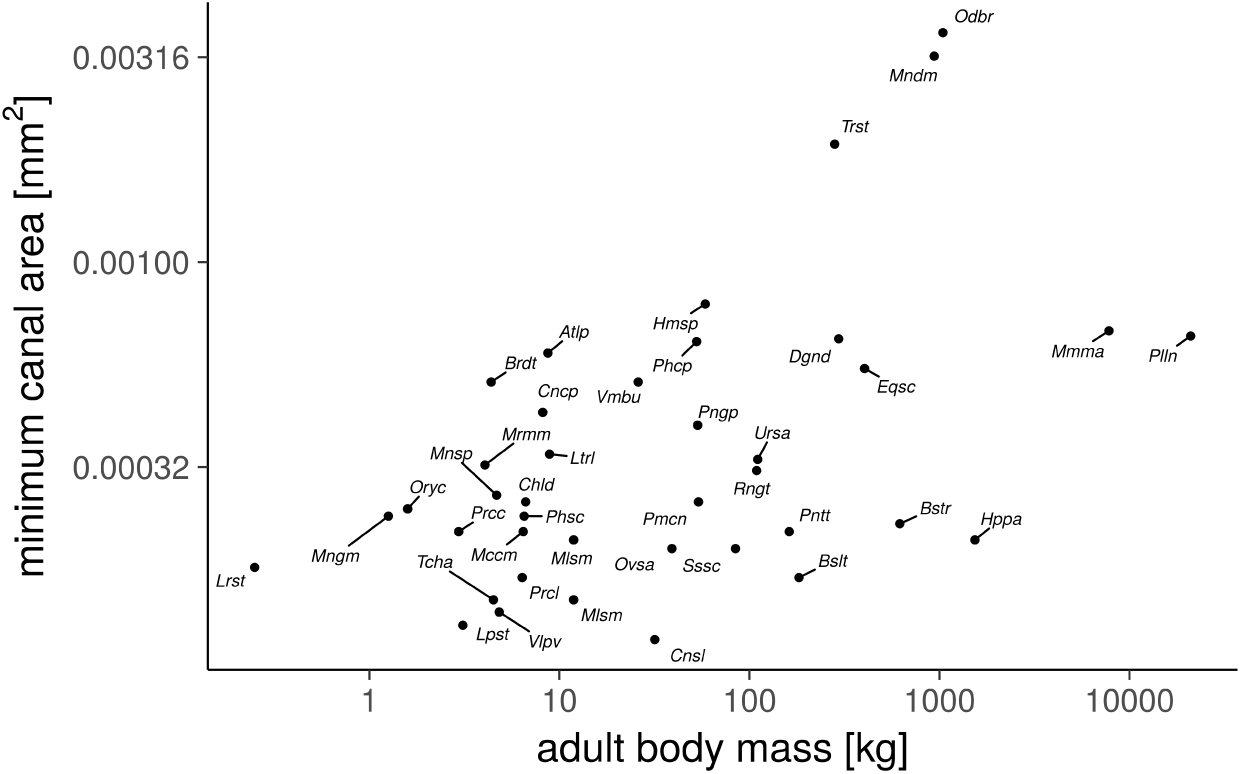
Log-log plot of per species minima of canal area and estimated body mass. Body mass relates to less than 30% of variation in minimum osteon area, canal area and infill area (Table 3), indicating that “narrow” secondary osteons are found throughout mammalian species.

**Table 3.** Summary of the scaling analysis for osteon area, canal area and infill area: slope estimate with lower and upper bounds (*b*,*b*^−^,*b*^+^), elevation estimate with lower and upper bounds (*a*,*a*^−^,*a*^+^), sample squared correlation coefficient (*R*^2^), and *p*-values for no correlation (*p*_*uncorrelated*_) and isometric scaling (*p*_*isometry*_) null hypotheses.

|  | osteon area |  |  |
| --- | --- | --- | --- |
|  | mean | minimum | maximum |
| $b$ | 0.23 | 0.23 | 0.27 |
| $b^-$ | 0.18 | 0.17 | 0.21 |
| $b^+$ | 0.28 | 0.31 | 0.33 |
| $a$ | 0.011 | 0.0047 | 0.018 |
| $a^-$ | 0.0087 | 0.0034 | 0.014 |
| $a^+$ | 0.014 | 0.0065 | 0.024 |
| $R^2$ | 0.54 | 0.14 | 0.53 |
| $P_{\text{uncorrelated}}$ | <0.005 | 0.017 | <0.005 |
| $P_{\text{isometry}}$ | <0.005 | <0.005 | <0.005 |
|  | canal area |  |  |
|  | mean | minimum | maximum |
| $b$ | 0.31 | 0.32 | 0.46 |
| $b^-$ | 0.24 | 0.25 | 0.36 |
| $b^+$ | 0.39 | 0.43 | 0.58 |
| $a$ | 0.00039 | 0.00011 | 0.00077 |
| $a^-$ | 0.00028 | 0.000071 | 0.00047 |
| $a^+$ | 0.00055 | 0.00016 | 0.0013 |
| $R^2$ | 0.45 | 0.27 | 0.46 |
| $P_{\text{uncorrelated}}$ | <0.005 | <0.005 | <0.005 |
| $P_{\text{isometry}}$ | <0.005 | <0.005 | <0.005 |
|  | infill area |  |  |
|  | mean | minimum | maximum |
| $b$ | 0.22 | 0.22 | 0.27 |
| $b^-$ | 0.18 | 0.17 | 0.21 |
| $b^+$ | 0.28 | 0.3 | 0.34 |
| $a$ | 0.01 | 0.0044 | 0.017 |
| $a^-$ | 0.0083 | 0.0032 | 0.013 |
| $a^+$ | 0.013 | 0.0061 | 0.023 |
| $R^2$ | 0.52 | 0.12 | 0.5 |
| $P_{\text{uncorrelated}}$ | <0.005 | 0.03 | <0.005 |
| $P_{\text{isometry}}$ | <0.005 | <0.005 | <0.005 |

### (b) Derived measurements

Shapiro-Wilk tests indicated that the logarithms of mean, maximum and median infill ratio were not normally distributed (p<0.005). We therefore performed non-parametric (Spearman’s rank) correlation tests on this data, which showed that mean, median and maximum infill ratio were not correlated with animal body mass (p>0.18 in all cases). Minimum infill ratio was not normally distributed (p<0.005, Shapiro-Wilk test) and a non-parametric test showed a very weak correlation with body mass (*p* < 0.05, *ρ*^2^ = 0.12). The significance of this correlation was lost when analyzing the data within separate groups of femoral and humeral specimens. Median infill ratio ranged from 84 to 99% (Figure 4). These results are summarized in Table 4.

**Table 4.** Summary of the scaling analysis for infill ratio and infill distance: slope estimate with lower and upper bounds (*b*,*b*^−^,*b*^+^), elevation estimate with lower and upper bounds (*a*,*a*^−^,*a*^+^), sample squared correlation coefficient, and *p*-values for no correlation (*p*_*uncorrelaied*_) and isometric scaling (*p*_*Sometry*_) null hypotheses. We report the median instead of the mean, and Spearman’s ρ, for the infill ratio, as the data were not normally distributed.

|  | infill ratio |  |  |
| --- | --- | --- | --- |
|  | median | minimum | maximum |
| **b** | -0.015 | -0.022 | -0.019 |
| **b<sup>-</sup>** | -0.02 | -0.029 | -0.026 |
| **b<sup>+</sup>** | -0.011 | -0.016 | -0.014 |
| **a** | 1 | 0.99 | 1 |
| **a<sup>-</sup>** | 0.97 | 0.96 | 0.99 |
| **a<sup>+</sup>** | 1 | 1 | 1 |
| **P<sub>shapiro-wilk</sub>** | **<0.005** | **<0.005** | **<0.005** |
| **$\rho^2$** | 0.0022 | 0.12 | 0.02 |
| **P<sub>non-param.</sub>** | 0.77 | 0.029 | 0.38 |
|  | infill distance |  |  |
|  | mean | minimum | maximum |
| **b** | 0.11 | 0.11 | 0.13 |
| **b<sup>-</sup>** | 0.089 | 0.084 | 0.097 |
| **b<sup>+</sup>** | 0.14 | 0.16 | 0.17 |
| **a** | 0.045 | 0.031 | 0.053 |
| **a<sup>-</sup>** | 0.04 | 0.027 | 0.046 |
| **a<sup>+</sup>** | 0.051 | 0.037 | 0.062 |
| **R<sup>2</sup>** | 0.44 | 0.073 | 0.33 |
| **P<sub>uncorrelated</sub>** | **<0.005** | 0.091 | **<0.005** |
| **P<sub>isometry</sub>** | **<0.005** | **<0.005** | **<0.005** |

Mean infill distance scaled with negative allometry (*b* = 0.121, *R*^2^ = 0.4, *p* < 0.001), indicating that while infill distance increases on average with species size in absolute terms, larger mammalian species have a relatively smaller mean infill distance (Table 4). We observed similar relationships for median and maximum infill distance. However, minimum infill distance is independent of species size, possibly due to the fact we included osteons at an early infilling stage. The overall maximum infill distance (found in specimen Bd 137, straight-tusked elephant) was 180 μm.

### (c) Phylogenetic signal

Scaling exponent estimates for phylogenetically independent contrasts of all variables of interest were slightly higher, but in all cases within the 95% confidence intervals obtained without any correction for phylogeny (Table 5), indicating minimal effect of phylogeny on the scaling relationships.

**Table 5.** Slope estimates, *R*^2^, and *p*-values for scale-invariant (*p*_*uncorrelated*_) and isometric scaling (*p*_*isometry*_) null hypotheses on the independent contrasts for our data.

| | $b_{independent\ contrasts}$ | $P_{uncorrelated}$ | $R^2$ | $p_{isometry}$ |
| --- | --- | --- | --- | --- |
| mean osteon area | 0.265 | <0.005 | 0.474 | <0.005 |
| minimum osteon area | 0.291 | 0.015 | 0.159 | <0.005 |
| maximum osteon area | 0.277 | <0.005 | 0.373 | <0.005 |
| mean canal area | 0.333 | <0.005 | 0.439 | <0.005 |
| minimum canal area | 0.368 | 0.005 | 0.204 | <0.005 |
| maximum canal area | 0.500 | <0.005 | 0.323 | <0.005 |
| mean infill area | 0.265 | <0.005 | 0.456 | <0.005 |
| minimum infill area | 0.286 | 0.020 | 0.145 | <0.005 |
| maximum infill area | 0.283 | <0.005 | 0.340 | <0.005 |
| mean infill ratio | −0.015 | 0.165 | 0.054 | <0.005 |
| minimum infill ratio | −0.023 | 0.049 | 0.106 | <0.005 |
| maximum infill ratio | −0.024 | 0.341 | 0.026 | <0.005 |
| mean infill distance | 0.129 | <0.005 | 0.369 | <0.005 |
| minimum infill distance | 0.153 | 0.068 | 0.092 | <0.005 |
| maximum infill distance | 0.142 | 0.009 | 0.180 | <0.005 |

## 4. Discussion

Our analysis shows that cross-sectional parameters of secondary osteons and their Haversian canals scale with negative allometry in the mammalian humerus and femur. Therefore, our key finding is that, where intra-cortical remodelling is present, larger mammals have larger osteonal and Haversian canal areas compared to smaller mammalian species in absolute terms, but smaller osteonal and Haversian canal areas in relative terms (Figure 6, Table 3). The distance between the canal border and the cement line also showed a negatively allometric relationship with animal size, whereas the ratio of infill area to osteon area does not scale (Table 4). Due to the large variation in sample cross-sectional area, unknown anatomical location and unclear boundaries of remodelled areas in some specimens, we did not feel comfortable reporting any osteon areal density (osteons/mm^2^) measurements, as they would not have been comparable between species. This was also not the focus of our study. Our images are in agreement with the general view that extensive remodelling [24] is only present in larger species. We did not find any secondary osteons in some medium-sized terrestrial species (Bd 8, *Macropus giganteus,* 33 kg; Bd 45, *Castor canadensis,* 18 kg; Bd 272, *Semnopithecus entellus,12* kg) and a large marine species (Bd 214, *Mirounga leonina,* 1600 kg), which might be due to the anatomical location or age of the specimen sampled.

**Figure 6.**
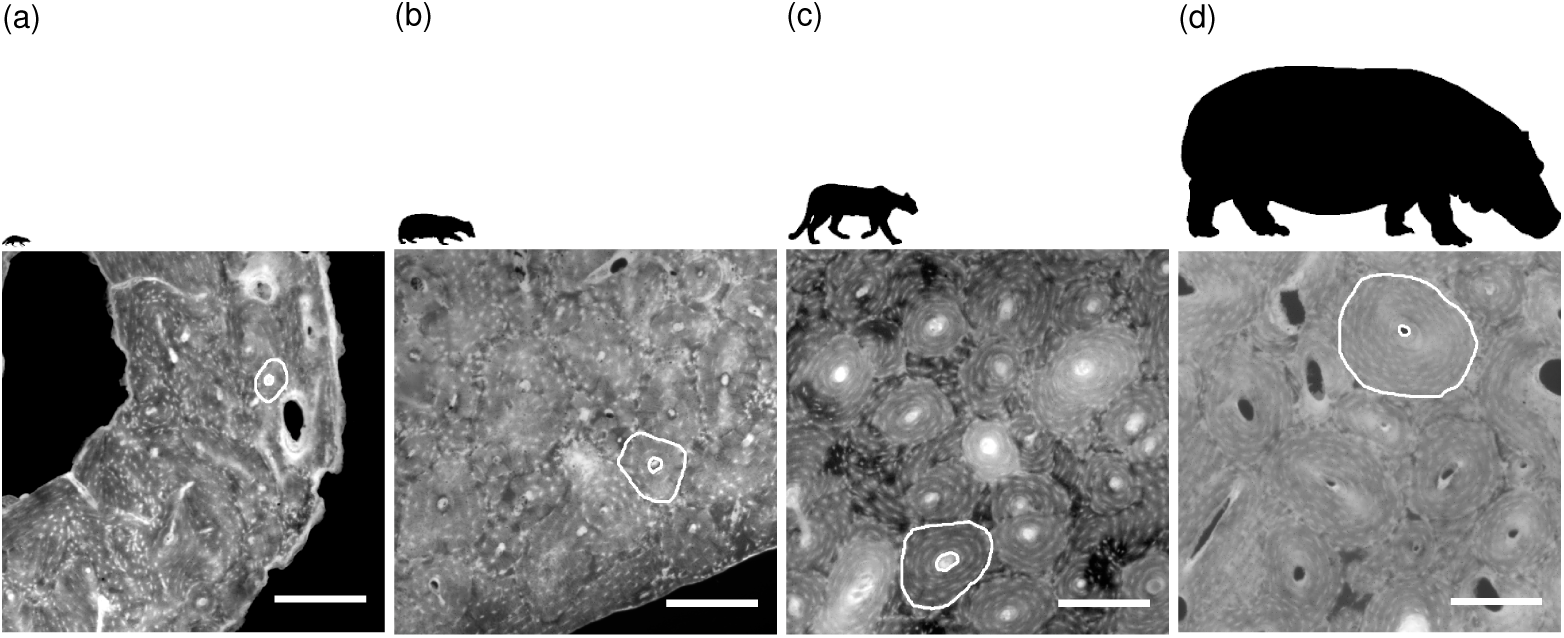
Close-ups of the bone cortex in four specimens with silhouettes reflecting the size differences above. (a) Banded mongoose, (b) European badger, (c) puma, (d) common hippopotamus (all scalebars 0.25 mm). Images ©The Royal College of Surgeons, and reproduced here with their kind permission.

### (a) Physiological and mechanical constraints on osteon dimensions

Covering an unprecedented range of mammalian species in which secondary osteons are found, the size-invariance of the infill ratio, and similar exponent estimates for osteon area and infill area (differing by less than 0.01) reported here confirm the results of previous studies showing a tight coupling between bone resorption and formation in intra-cortical bone remodelling. Qiu and coworkers [68] demonstrated a strong (r=0.97) quadratic relationship between osteon perimeter and infill area in young and elderly humans. Another study [69] showed strong correlations (r>0.95) between osteon area and infill area for osteons of varying sizes from several species. Thus, seemingly, a similar percentage of resorbed bone is filled back in, independent of body size, suggesting that it is crucial for all mammalian limb bones to keep their overall porosity at a similar level, since porosity is an important determinant of bone strength [70].

Sample squared correlation coefficients are lower for minimum than for maximum and mean osteon, canal and infill area. Erythrocyte dimensions may form a lower limit for the cross-sectional area of remote blood vessels such as Haversian canals and do not vary with species size in mammals [71]. Idealising the minimum canal areas we measured into circles resulted in diameters of approximately 10 μm, which are close to the diameter of an erythrocyte (approx. 8 μm, [72]), supporting this notion. Therefore, occasionally, osteons of large species are narrow relative to the entire, inter-species range we report in the present study (and the erythrocytes still fit through the capillaries), but wide osteons do not occur in small species.

On the other hand, upper limits for osteon and canal area may differ for species of varying size: The maximum distance from a blood vessel at which osteocytes remain viable has been reported to be 230 μm [15]. This is close to the maximum infill distance we observed in the larger species of our sample and suggests that maintaining osteocyte viability, combined with the previously mentioned size-invariant infill ratio, constitutes a limiting factor for osteon area in large species. For smaller species, however, this limit is never reached, suggesting that a different mechanism dominates the size of osteons in these animals. The cortical thickness of the specimen with the smallest estimated body mass and evidence of osteons in our sample (Red slender loris, *Loris tardigradus)* ranges between 600 and 850 μm, which is of comparable scale to the estimated diameter of the largest osteon we found overall (580 μm). A resorption area of that size in as thin a cortex is likely to have deleterious consequences for the bone [73]. Creating as large osteons as possible, if an organism can afford the temporary increase in porosity due to a large resorption cavity, may be advantageous from a mechanical point of view after infilling has completed. Larger (and more numerous) osteons have been associated with increased toughness [74–77]. It has more recently been suggested that larger osteons [78] are geometrically more advantageous for resisting bending and compression loads. Studies treating bone as a quasi-brittle material suggest that bone strength and toughness improve with increases in the ratio between dominant inhomogeneity size (i.e. in our case, the typical size of microscopic features such as osteons) and the structural scale (i.e. the bone organ size) [79–81]. Therefore, larger osteons, to the extent to which they do not preclude osteocyte viability, in large species may be explained as tissue-level geometrical adaptations to offset mechanical disadvantages that come with increased size, conceptually similar to adaptations at the organ level described in previous studies such as decreasing bending moments with more upright postures [82], increased limb bone robustness [83], cortical thickness and infilling with trabeculae [84], and adapting number and thickness of trabeculae [21].

The fact that phylogeny had little effect on our findings is unsurprising, given the diversity of species, clades and ecologies that we sampled from. This also reinforces the notion that our results likely reflect overall bio-physical limits within which the intracortical bone remodelling process operates, and not clade- or species-specific adaptations.

### (b) Limitations

The high variability of our data reflects the multitude of species- and specimen-specific characteristics that will affect bone remodelling, including age, sex, locomotor style, life history, onset of secondary remodelling during ontogeny, environmental influences and more. Specimen age was unknown, but we expect most specimens to be of young adult age, as life expectancy in 19th century zoos was low ([85], as cited in [86]). There is some debate about the relationship between age and osteon size in humans: While most studies support an inverse relationship (narrower osteons in older people) [87,88], some studies show no effect. Osteon size was not correlated with age in macaques [31]. The effect size of age in interspecies comparisons of secondary osteon dimensions (if there is one) is however likely to be smaller in magnitude to the relationship of secondary osteons with mass that we show here. Similarly, we could not take sex into account in our analysis, since it was unknown for all our specimens.

Because stylopodal dimensions also scale with body size [19], it seems reasonable to speculate that bone size, and not body size, determines osteon dimensions [89]. However, data for calcanei and femora from equine samples, and metapodials and femora from American black bears, suggests that secondary osteon dimensions are similar across different anatomical sites and across bones of varying size within the same species [69]. This makes it appear more likely that osteon dimensions relate to adult body mass, as our data show, and not to bone size. To distinguish the contributions to osteon size of bone organ size and whole body mass in more detail, a further study, similar to ours, comparing osteon dimensions in bones of the same size across species, and in bones of different sizes within species, would be required.

More insights would be gained if we were to measure the secondary remodelling process in three dimensions. However, while we are now able to image resorption cavities effectively using X-ray microtomography [90], for now, the characterization of cement sheaths in 3D involves considerable manual effort [4] and could not have been completed within the time-scale of this study.

Using a historical collection of slides presented some further limitations. To an extent, knowledge of exact anatomical location, precise taxon as well as specimen age and provenance was sacrificed in order to sample a large species and size range. We feel this was important, as we would have been prone to concluding that there was no scaling relationship if we had used a smaller number of different species that are easily accessible, given that (for example) humans have relatively wide, and cattle have relatively narrow, osteons for their size. The use of a historical collection additionally avoided the time, effort and difficulties required to obtain and generate an equivalent collection to a modern standard.

Fluorochrome labelling could have helped validate our results. However, this would have constituted only a partial validation: it would have confirmed whether we characterized the secondary osteons formed during the labelling period correctly (i.e. true positives), but would likely have produced a number of false negatives, namely intact secondary osteons formed prior to fluorochrome injection. Using solely labelled osteons would have reduced the number of osteons sampled, increased the bias towards osteons which had not completed infilling at time of death or had possibly differing infilling rates, and could have therefore confounded our results. Moreover, obtaining labelled material from a large enough range of species would have posed considerable difficulties.

### (c) Conclusion

In summary, the images of historically important specimens we produced are available (See data accessibility statement below) to researchers for use in future comparative histological studies. The measurements of secondary osteon and Haversian canal area we presented here have the highest number of mammalian species with the most diversity in clades and the broadest range in size to date. We performed a comprehensive scaling analysis of quantities describing the intra-cortical remodelling process across species, which showed negative allometry for osteon, canal and infill area as well as infill distance, while the infill ratio was size-independent. Our results indicate that minimum osteon dimensions correlate only very weakly with animal body mass and might relate to erythrocyte dimensions. In contrast, the upper limits to osteon dimensions display a strong relationship with adult body size: osteons in small species may be restricted to sizes avoiding fatal temporary stress increases around too large resorption cavities, while osteon size in large species may be dictated by the ability to maintain osteocyte viability.

#### Data Accessibility

Full-resolution images can be obtained from RCS for research purposes upon request (Low-resolution versions can be found on the RCS online catalogue SurgiCat by searching for the reference numbers listed in Tables 1 and 2). Segmentations of the images, and the raw data of the histomorphometrical measurements are available online. The code used to obtain our results is available on github [66,67].

#### Authors’ Contributions

AF imaged the specimens, hand-traced the secondary osteons, wrote the code for image and statistical analysis, carried out the statistical analyses and drafted the manuscript; CP, HC and MC searched the RCS collection and archives and prepared the slides for imaging; CP, HC, MC and AF drafted paragraphs about the history and condition of the Quekett slides and reviewed the manuscript; MD conceived of and coordinated the study and helped draft the manuscript; JRH conceived of the study, helped draft the manuscript and gave advice on phylogenetic correction. All authors gave final approval for submission and publication.

#### Funding

AF’s PhD project is funded by The Royal Veterinary College and Foster+Partners. The study presented here was a collaborative effort between RCS and RVC.

## Acknowledgements

The authors thank all the RCS staff and the Quekett Microscopy Club for providing access to the Quekett collection. AF is grateful to Andrew Cuff for help with phylogenetic independent contrasts, to Andrew Pitsillides, Inês Perpétuo and Jim Usherwood for helpful discussions, and to Richard Domander for reviewing the code.

